# FEELnc: A tool for Long non-coding RNAs annotation and its application to the dog transcriptome

**DOI:** 10.1101/064436

**Authors:** V Wucher, F Legeai, B Hédan, G Rizk, L Lagoutte, T Leeb, V Jagannathan, E Cadieu, A David, H Lohi, S Cirera, M Fredholm, N Botherel, P Leegwater, C Le Béguec, H Fieten, C Johansson, J Johnsson, LUPA consortium, J Alifoldi, C André, K Lindblad-Toh, C Hitte, T Derrien

## Abstract

Whole transcriptome sequencing (RNA-seq) has become a standard for cataloguing and monitoring RNA populations. Among the plethora of reconstructed transcripts, one of the main bottlenecks consists in correctly identifying the different classes of RNAs, particularly those that will be translated (mRNAs) from the class of long non-coding RNAs (lncRNAs). Here, we present FEELnc (FlExible Extraction of LncRNAs), an alignment-free program which accurately annotates lncRNAs based on a Random Forest model trained with general features such as multi *k*-mer frequencies and relaxed open reading frames. Benchmarking versus five state-of-art tools shows that FEELnc achieves similar or better classification performance on GENCODE and NONCODE datasets. The program also provides several specific modules that enable to fine-tune classification accuracy, to formalize the annotation of lncRNA classes and to annotate lncRNAs even in the absence of training set of noncoding RNAs. We used FEELnc on a real dataset comprising 20 new canine RNA-seq samples produced in the frame of the European LUPA consortium to expand the canine genome annotation and classified 10,374 novel lncRNAs and 58,640 new mRNA transcripts. FEELnc represents a standardized protocol for identifying and annotating lncRNAs and is freely accessible at https://github.com/tderrien/FEELnc.

## INTRODUCTION

The development of high-throughput RNA sequencing (RNA-seq) has allowed the identification of many RNA species in different organisms such as in mammals (1–3), insects (4, 5) or plants (6, 7). Particularly, whole transcriptome sequencing shed light on the pervasive transcription of the genomes with messenger RNAs (mRNAs) only representing a small fraction of the genome, outnumbered by a vast repertoire of small and long non-coding RNAs (lncRNAs). LncRNAs, basically defined as transcripts longer than 200 nucleotides and without any protein-coding capabilities, have been involved in many aspects of normal and pathological cells. From the pioneer discovery of the *Xist* lncRNA involved in X chromosome inactivation in placental females (8) to the more recent links between lncRNAs and cancers (9,10), lncRNAs have emerged as key actors of the cell machinery with diverse modes of action such as gene expression regulation, control of translation or imprinting.

Following RNA sequencing, the computational reconstruction of transcripts models either by genome-guided (11, 12) or *de novo* assembly (13) usually produces tens of thousands of known and novel transcript models. Among this wealth of assembled transcripts, it remains crucial to annotate the different classes of RNAs and especially to distinguish protein coding from non-coding RNAs. To this aim, several bioinformatics tools have been developed in order to compute a coding potential score (thereafter termed CPS) used to discriminate the coding status of the RNA gene models. Broadly, they can be divided into programs using sequence alignments either between species (14) or alignments to protein databases (15) and alignment-free software (16–18). The alignment-dependent methods, although very specific in term of performance, are often very time and resource consuming. For instance, the PhyloCSF program requires a multispecies sequence alignments to predict the likelihood of a sequence to be a conserved protein-coding based on the evolution of the codon substitution frequencies (14). It is thus dependent on the quality of the input alignments and may also be biased towards misclassifying species-specific or lowly conserved coding and non-coding transcripts (19). On the contrary, the alignment-free methods compute a CPS only depending on intrinsic features of the input RNA sequences. One of the main features is given by the length of the longest Open Reading Frame (ORF) (20, 21) where usually a transcript harbouring a long ORF will most likely be translated into proteins. However, the definition of the longest ORF can vary between programs especially when it involves the strict inclusion of either or both the start and the stop codons. This is particularly important to model since some transcripts from reference annotations and/or newly assembled transcripts are not full-length. For instance, the proportion of protein-coding transcripts in the human EnsEMBL (v83) annotation (22) lacking a start codon or a stop codon is 10% (7,677/79,901) and ~25% (16,649/79,901), respectively. A complementary feature to discriminate mRNAs from noncoding RNAs is given by the relative frequency of oligonucleotides or *k*-mer (where *k* designs the size of the oligonucleotide). Some tools already used *k*-mer frequencies but are often limited to one or small *k*-mer (generally *k* <= 6) whereas longer *k*-mer could help resolving ambiguities by taking into account lncRNAs-specific repeats or spatial information for instance (23, 24). Finally, common to all methods is the lack of an explicit modeling and cutoffs definition for “non-model” organisms (25) for which it could be crucial to train the programs with species-specific data and to automatically derive a CPS cutoff which provide better discriminative power.

Here we present FEELnc, for FlExible Extraction of LncRNAs, a new tool to annotate lncRNAs from RNA-seq assembled transcripts. FEELnc is an all-in-one solution from the filtering of non-lncRNA-like transcript models, to the computation of a coding potential score and the formalization of the definition of the lncRNA classes. As an alignment-free method, the program employs Random Forest (26) to classify lncRNA and mRNAs based on a relaxed definition of ORFs and a very fast analysis of small and large *k*-mer frequencies (from k = 1 to 12). We benchmarked FEELnc versus five existing programs (CNCI, CPC, CPAT, PhyloCSF, PLEK) using known set of lncRNAs annotated in multiple organisms (GENCODE for human and mouse (27) and NONCODE for other species (28)), and showed that FEELnc performance metrics outperformed or are similar to state-of-the-art programs. We developed FEELnc to be used on “non-model” organisms for which no set of lncRNAs is available by deriving species-specific lncRNA models which are shuffled from mRNA sequences and automatically computing the CPS cutoff that maximizes classification performances. FEELnc also allows user to provide its own specificity thresholds in order to annotate high-confidence sets of lncRNAs and mRNAs and to define a class of ambiguous transcript status. Finally, we produced 20 RNA sequencing dataset as part of the LUPA consortium (29) sampled from 16 different canine tissues and applied FEELnc to annotate 10,374 new lncRNAs and 58,640 novel mRNA isoforms. We also classified the new lncRNAs into 5,033 intergenic (lincRNAs) and 5,341 genic sense or antisense lncRNAs based on the FEELnc classifier module. The number of lncRNA transcripts detected by our data considerably expands the canine annotation providing an extended resource which will help to deciphering genotype to phenotype relationships (30).

## MATERIAL AND METHODS

### 1. Datasets

For the sake of reproducibility, all datasets and scripts made to generate the benchmark files are available in supplemental materials.

Human long non-coding and protein-coding genes were downloaded from the manually curated GENCODE version 24 (EnsEMBL v83 on the GRCh38 human genome assembly) with the following biotypes “ *lincRNA”* or *“antisense”* and *“protein_coding”,* respectively. From this initial dataset, we extracted 10,000 mRNAs and 10,000 lncRNAs that were divided into two files for the learning and the testing steps (denoted HL and HT, respectively), each of these files contains 5,000 mRNAs and 5,000 lncRNAs. Importantly, only one transcript per locus was extracted for all biotypes in order not to bias by introducing two isoforms of the same gene in both the HL and HT sets. For mouse, we used the GENCODE version M4 (EnsEMBL v79) and derived the learning and testing files (denoted ML and MT) as for human. Due to the lower number of GENCODE lncRNAs annotated in mouse compared to human, each file contains ~2,000 lncRNAs and 5,000 mRNAs. For “non-model organisms”, lncRNAs belonging to the antisense and intergenic classes (NONCODE codes 1000 and 0001, respectively) were downloaded from the latest version of the NONCODE database (NONCODE 2016) (28) while mRNAs were retrieved from the EnsEMBL database (v84). A summary of the number of mRNAs/lncRNAs per species is available (Supplementary Table 1).

Whole transcriptome sequencing of dog samples (n = 20) were generated in the frame of the LUPA consortium. The biological samples were obtained from the “Cani-DNA_CRB” (http://dog-genetics.genouest.org) and XXX biobanks. The dog owners consented to the use of data for research purposes anonymously. The sequencing libraries were constructed using paired-end stranded polyA selection and sequenced by HiSeq-2000 Illumina technology. The 20 samples correspond to 16 unique tissues and 7 breeds (Supplementary Table 2) with about 50 Millions reads per sample. The RNASeq data is available in the short read archive (SRA) under NCBI bioproject PRJNA327075 and SRA accession SRP077559.

### 2. FEELnc coding potential predictors

Among the three FEELnc modules, the second one aims at computing a coding potential score given the assembled sequences following transcriptome reconstruction. To deal with the incompleteness of ORF annotation where both the reference and the reconstructed transcripts may not be full-length, FEELnc can compute five types of ORF from the stricter “type 0” which corresponds to the longest ORF having both a start and stop codons, to the more relaxed “type 4” that is the whole input RNA sequence (see Supplementary Material for detailed description of the ORF types). Because the size of the protein-coding ORF is generally correlated with the length of the input RNA sequence, we used the **ORF coverage**, i.e. the proportion of the transcript size covered by the ORF, as the first predictor to discriminate mRNA/lncRNAs in the FEELnc model.

The second predictor of FEELnc relies on the computation of the **multi *k*-mer frequencies** between mRNAs and lncRNAs. Biases in nucleotide frequencies and codon usage have already been described in the literature as important discriminative features between coding and noncoding RNAs (21, 31). Within the framework of FEELnc, we developed an extremely fast and exact *k*-mer counter called KIS (for *k*-mer In Short) that relies on the open-source GATB library (32). For example, KIS can compute all 6-mers (hexamers) and 12-mers of the human GRCh38 genome assembly (n~3,1 billions *k*-mers) in ~2min and 2m51, respectively. Due to KIS high speed, a major contribution of this work was to be able to combine different list of *k*-mers including longer *k*-mers in order to better discriminate lncRNAs from mRNAs. In more details, we assigned a score for each *k*-mer based on the occurrence of all *k*-mers of size *k* in each predicted ORF sequences for both lncRNAs and mRNAs. The score 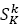 similar to the one used in Claverie et al. in (33), is computed for each specific *k*-mer *K* of size *k* as follow:

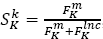, with 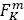 and 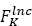 the observed frequency of *K* in mRNA ORFs and in lncRNA ORFs for the two training sets, respectively. Once the *k*-mer model is made for a size *k*, i.e. all 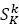 have been computed, FEELnc associates a *k*-mer value 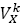 for a sequence *X* as follow:

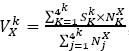, with 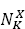 the occurrence of *K* in sequence *X* and 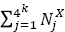 the total number of *k*-mer of size *k*. Using this scoring method, one *k*-mer score is associated to each sequence for each *k*-mer size selected in the model. Note that the scanning of the predicted ORF sequences is done using a step size of 3 for the all sequences of the testing file and the mRNA learning file and 1 for the lncRNA learning file.

FEELnc coding potential also uses the total **RNA sequence length** as a predictor of the model since lncRNAs are known to be significantly shorter than mRNAs (34, 35). For illustration purpose, a distribution of the FEELnc predictor scores with “type 3” ORF and multi *k*-mer scores with *k* in {1,2,3,6,9,12} is given in Supplementary Fig.1 regarding the 5,000 mRNAs and 5,000 lncRNAs of the HL dataset.

### 3. Random Forest classification and optimized coding potential cut-offs

The aforementioned predictor scores are incorporated into a machine learning method – here Random Forest (RF) (26) - that computes a Coding Potential Score (CPS) for each input training transcripts. As also shown by others (36), RF often outperforms other machine learning techniques especially due to the random sampling of features to build the ensemble of trees. In addition, our RF model could deal with imbalanced training set by down-sampling the majority class (most likely mRNAs in many organisms). In fact, the CPS in our RF model corresponds to the proportion of all trees (500 trees by default) which “vote” for the input sequence to be coding or non-coding. A proportion close to 0 will indicate a non-coding RNA and close to 1 for mRNA.

To define an optimal CPS cut-off, FEELnc automatically extracts the CPS that maximizes both sensitivity (Sn) and specificity (Sp) performance (see performance section) based on a 10-fold cross-validation. Using the ROCR package (37), FEELnc provides user with a two-graph ROC curve in order to display the performances of the model and to visualize the optimal CPS (Fig. 1A).

**Figure 1:**
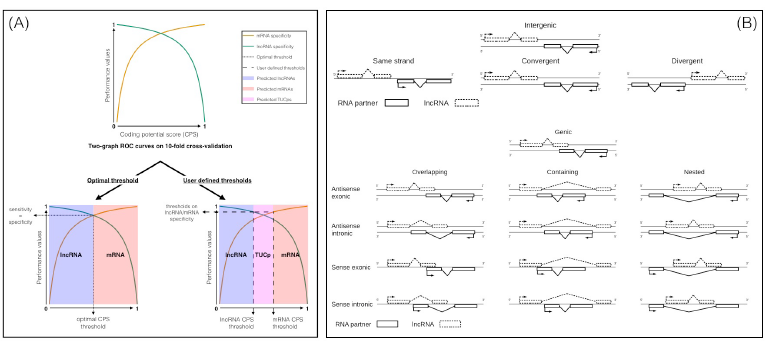
A) Two graph ROC curves for automatic detection of optimized CPS threshold and user specificity thresholds, the latter defines two conservative sets of lncRNAs and mRNAs and a class of ambiguous biotypes transcripts termed TUCp (Transcript of Unknown Coding potential). B) Sub classification types of intergenic transcripts/lncRNA interactions and sub classification types of genic transcripts/lncRNA interactions.

Even if this approach aims at providing highest performances, it could sometimes misclassify transcripts whose CPS is closed to the optimized threshold (38). To take this into account, FEELnc allows fixing two minimal specificity cut-offs for lncRNAs and mRNAs (this approach is termed “2-cutoffs”). This naturally leads to the annotation of two high-confident classes of lncRNAs and mRNAs and also the definition of a third class of ambiguous transcripts (i.e transcripts for which the CPS is in-between the 2 cutoffs) that are named TUCp (38) for Transcripts of Unknown Coding potential (Fig. 1A).

### 4. FEELnc without long non-coding training set

One issue when using machine learning algorithms is the requirement to have both mRNA and lncRNA learning sets to train the model. While the former is often available in most of the organisms, the latter is still in its infancy for some species and especially for non-model organisms (25). To model non-coding RNAs in the absence of a true set of lncRNAs, we assessed three strategies called “*intergenic*”, “*shuffle*” and “*cross-species*”. The first one consists in extracting random intergenic sequences of length *L* (*L* given by the distribution of the mRNA sizes) as the non-coding training set. The second employs a different approach by shuffling mRNA sequences from the reference annotation using the Ushuffle program (39) while preserving the *k*-mer frequency given a fixed *k* length of the input sequences. Note that with increasing length of *k*, it is possible that Ushuffle could not permute some input sequences because of the constraint to preserve *k*-mer frequency. Finally, the strategy *cross-species* makes use of lncRNA sets annotated in other species to extract noncoding predictors and train the RF model. For the latter strategy, we used the NONCODE lncRNA catalogues in 13 species and train FEELnc model using human mRNAs and NONCODE species-specific lncRNAs. For all strategies, we assessed the performance on the HT datasets.

### 5. FEELnc classifier

Given a known reference annotation, it is essential to classify newly annotated lncRNAs based on their closest annotated transcripts. It will potentially guide biologists towards functions and relationships between lncRNAs and its annotated partners (lncRNA/mRNA pairs for instance). For this purpose, the FEELnc classifier module employs a sliding window strategy (whose length is fixed by user) that scans within and around each lncRNA all overlapping transcripts from the reference. Not only FEELnc classifier annotates intergenic lncRNA (lincRNAs) or antisense but the module also formalizes the definition of lncRNA subclasses with respect to annotated transcripts (Fig. 1B). First, these rules involve the *direction* (sense or antisense) and the *type* of interactions (genic or intergenic). Then, within each *type* of interaction, a *subtype* level allows to narrow down the classification (e.g. divergent for lincRNAs or containing for genic lncRNAs). Finally, a *location* level is added which informs on the position of the lncRNAs with respect to the annotated transcripts (e.g. upstream for lincRNA or exonic for genic lncRNAs). In addition, the FEELnc_classifier_ can be used with all transcript biotypes (e.g. short ncRNAs such as snoRNAs) from a reference annotation and therefore is capable of annotating lncRNAs host gene for short RNAs (40) (hence the “*genic sense exonic*” class).

Because one lncRNA could belong to different classes depending on which reference transcript is considered, our approach reports all interactions within the defined widow and defines a best partner transcript using the following priorities: a lncRNA overlapping a reference transcript exon will be prioritized over intronic (and over containing) for genic lncRNAs, while the nearest reference transcript will be selected for lincRNAs.

### 6. Performances

We evaluated the performances of FEELnc and five other programs: CNCI (version 2 Feb 28, 2014), CPC (version 0.9-r2), CPAT (version 1.2.1), PhyloCSF (version 20121028-exe) and PLEK (version 1.2) by computing classical performance metrics:

– Sensitivity (Sn) or True Positive Rate = 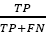,
– Specificity (Sp) or True Negative Rate = 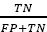,
– Precision (Prec) or Positive Predicted Value = 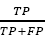,
– Accuracy (Acc) = 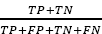,

with TP: True Positive, TN: True Negative, FP: False Positive and FN: False Negative.

In addition, we used two complementary metrics that could be considered as more global values of performance.

– F1-score = 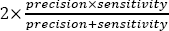, which is a statistic measuring the harmonic mean of precision and sensitivity.
– MCC (Matthews Correlation Coefficient) = 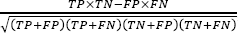, which is particularly useful when the two classes are of very different sizes (often the case between mRNA and lncRNA in non-model organisms) and which could be seen a correlation coefficient between trained and tested datasets.

For each performance metric, we considered lncRNAs as the negative class and mRNAs as the positive class. As a binary classification, the mRNA specificity corresponds to the lncRNA sensitivity (and conversely). For the CPAT and PLEK programs, which allow training their models, we used species-specific set of mRNAs/lncRNAs for training (called CPAT_train_ and PLEK_train_, denoted as CPAT_train and PLEK_train in Figures and Tables). For CPAT_train_, we referred to the optimal CPS cut-off mentioned on the website to discriminate between coding and non-coding RNAs (18). A detailed description of the command lines used to run each program is given in Supplementary Materials.

## RESULTS

### 1. FEELnc modules to annotate lncRNAs

Starting from assembled transcripts and a reference annotation, the FEELnc pipeline is composed of three independent modules to classify and annotate lncRNAs (Fig.2). The first module (FEELncfilter) aims at identifying and removing non-lncRNA transcripts from the reconstructed transcript models. To achieve this goal, FEELnc removes every assembled transcript that overlaps in sense any exon of the reference annotation. Importantly, FEELnc enable to parameterize the percentage of overlap and the transcript biotype (e.g *“protein coding”* or *“pseudogene”)* from the reference annotation. Indeed, to identify lncRNAs, transcripts matching protein-coding exons shall be removed as they likely indicate novel mRNA isoforms. Consequently, novel transcripts overlapping other short ncRNAs for instance will be kept as these may correspond to long non-coding RNAs host genes for small RNAs for instance (41). FEELnc_filter_ filters out short transcripts (default 200 nt) and can deal with single-exon transcripts depending on whether the protocol used to construct libraries is stranded or not. Remaining transcripts are thus candidates to be new lncRNAs or mRNAs.

**Figure 2:**
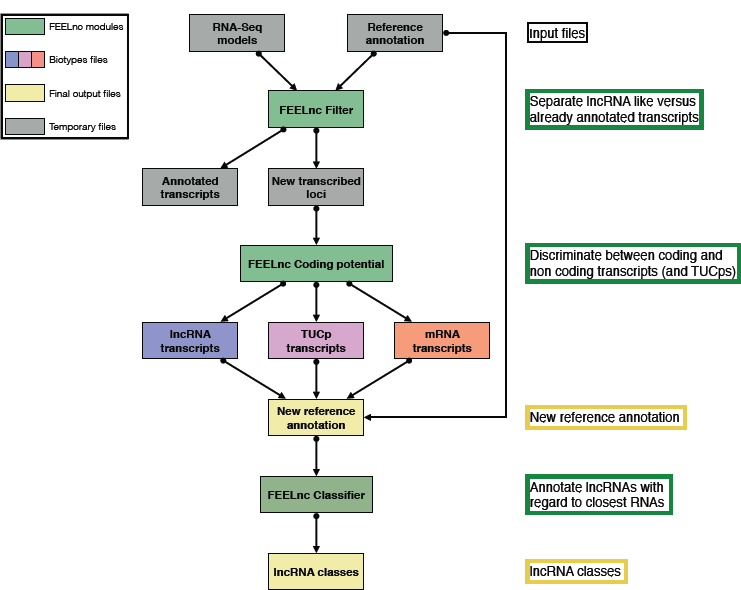
General overview of the FEELnc pipeline. The FEELnc filter module identifies newly assembled RNA-seq transcripts and removes non-lncRNA transcripts. The FEELnc coding potential module computes a coding potential score (CPS) and automatically defines the optimal CPS score cut-off to discriminate lncRNAs versus mRNAs (and eventually TUCPs). The FEELnc classifier annotates lncRNAs classes based on RNAs from the reference annotation.

The second module (FEELnc_codpot_) computes a coding potential score for every candidate transcripts based on a RF model trained with several predictors such as ORF coverage, multi *k*-mer frequencies and RNA sizes (see Methods). Using the “gold-standard” GENCODE human learning set (HL) for training, we evaluated the performances of FEELnc on 5,000 lncRNAs and 5,000 mRNAs from the test dataset with respect to the five ORF types and multi *k*-mer combinations. We observed that “type 1” and “type 3” ORF models, which extract the longest ORF even in the absence of stop codon, consistently display better achievements (mean MCC = 0.816) than “type 0” and “type 2” ORFs (mean MCC = 0.67 and 0.68, respectively) whatever the combination of *k*-mer is considered (Supplementary Fig.2). In addition, the multi *k*-mer strategy improves the performance of the program with a MCC performance starting at 0.80 when only using *6*-mers but reaching 0.85 with a combination of *{1,2,3,6,9,12}-*mers (with a fixed ORF type 3) (Supplementary Fig.2).

In addition to measuring performance, we conducted several evaluations to assess the robustness of FEELnc predictions. First, we showed that FEELnc is not biased by unbalanced or low numbers of transcripts in the input training set with sensitivity and specificity values higher than 0.9 using only 400 mRNAs and lncRNAs (Supplementary Fig.3). Second, FEELnc performs similarly or better than other methods to classify small or long mRNAs and lncRNAs (Supplementary Fig.4). Third, in order to model incompletes RNA reconstruction, we removed 10%, 25% and 50% from the 5’ of each sequence from HT data set. Even if FEELnc_codpot_ performance decreased with increasing degradation percentage (MCCs = 0.827, 0.749 and 0.53 for 10%, 25% and 50%, respectively), it performed better than other tested methods (Supplementary Fig.5). Fourth, we were able to replicate the performance with mouse GENCODE dataset ML and MT (see materials) composed of 5,000 mRNAs and 2,000 lncRNAs where FEELnc achieves 0.938 in sensitivity, 0.941 in specificity and a MCC of 0.856. Finally, among the manually curated list of 35 “well-characterized” lncRNAs from (42), FEELnc correctly classifies 33 (95%) of them as non-coding. The two discordant lncRNAs are *PWRN1* which has a CPS just above the cutoff (0.374 versus 0.372) and more surprisingly, the lncRNAs *FIRRE* having the highest CPS = 0.58 (the median CPS of the 35 lncRNA is 0.088). The high protein-coding score for *FIRRE* lncRNA can be explained by a large and complete ORF (546 nt i.e. 58% of the total RNA sequence) and a high level of similarity with the *FAM195A* protein-coding gene. Using the 2-cutoff strategy (see Method) with specificity cutoffs fixed at 0.95 for both mRNA and lncRNA biotypes, FEELnc classifies *PWRN1* and nine other lncRNA as ambiguous (TUCps) but still predicts *FIRRE* as the only protein-coding transcript.

The third FEELnc module (FEELnc_classifier_) formalizes the annotation of lncRNAs based on neighbouring genes in order to predict lncRNA functions and RNAs partners (see classes in methods). To illustrate the outcome of the FEELnc_classifier_, we applied it on the human EnsEMBL v83 annotation composed of 24,659 lncRNAs (“ *lincRNA”* and *“antisense”* biotypes) and 79,901 mRNA transcripts *(“protein coding”* biotype). For instance, FEELnc_classifier_ annotates 5,544 *“lncRNAs intergenic antisense upstream”* among which 40% (n = 2,234) are less than 5kb from their mRNA partner transcription start sites (TSS). This class directly pinpoints to lincRNAs potentially sharing a bidirectional promoter with its mRNA partner (43). At the opposite, 408 lincRNAs located less than 5kb from the mRNA, belongs to the *“sense intergenic upstream”* class and may correspond to dubious lncRNAs that are actually 5’UTR extensions of the neighboring protein-coding RNAs. FEELnc_classifier_ also annotates 5,006 lncRNAs in the *“antisense exonic”* class as potential candidates for complementary interactions with the mRNA transcribed in opposite direction (44, 45).

### 2. Benchmarking FEELnc in comparison with existing tools

We next compared the performance of FEELnc with five state-of-the-art programs either alignment-free (CPAT, CNCI and PLEK) or alignment-based (PhyloCSF and CPC). As for FEELnc, CPAT and PLEK allow building their models using user-specified learning dataset. Hence, we used the balanced HL dataset (5,000 lncRNAs and 5,000 mRNAs) to construct the models for these two programs denoted CPAT_train_ and PLEK_train_. Contrary to FEELnc, PLEK and CPAT required to a priori extract the CDS of the mRNA input file to learn the coding parameters. We also used default pre-built models for PLEK and CPAT although some of the transcripts from the human GENCODE dataset test file (HT) could have been used for building these models. For all programs, performance metrics were calculated according to the human GENCODE dataset (HT). It showed that FEELnc had the highest classification power (AUC value 0.97) compared to the others tools as illustrated by the ROC curve (Fig.3A).

**Figure 3:**
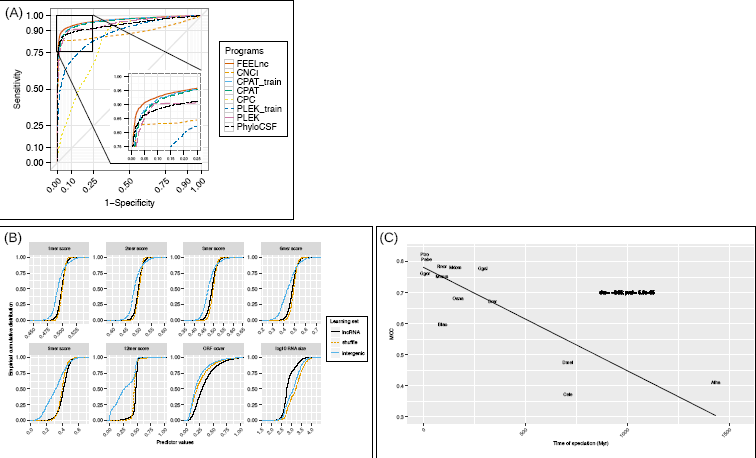
A) ROC curve analysis of FEELnc versus ncRNA annotation programs based on GENCODE human dataset (HT). B) Empirical cumulative distribution of FEELnc_codpot_ feature scores with the true set of human lncRNAs (lncRNA) in comparison with the “*shuffle*” and “*intergenic*” approaches. C) FEELnc_codpot_ MCC values tested on human set and trained using human mRNAs and species-specifc NONCODE lncRNAs. In x-axis is represented the time of speciation between human and NONCODE species as given in (57). Species abbreviations are the following: Atha: Arabidopsis; Btau: Cow; Cele: Nematode; Dmel: Fly; Drer: Zebra; Ggal: Chicken; Ggor: Gorilla; Hsap: Human; Mdom: Opossum; Mmul: Rhesus; Mmus: Mouse; Oana: Platypus; Pabe: Orangutan; Ptro: Chimpanzee; Rnor: Rat.

Accordingly, FEELnc displays the highest sensitivity (0.923) and secondly ranked specificity (0.915) values among all tools while PLEK displays the highest specificity (0.985) and precision (0.981). In general, alignment-based methods have lower classification metrics than alignment-free programs and it appears that most of the CPC misclassified lncRNAs are lincRNAs (37%) versus antisense lncRNAs (10%). Finally, FEELnc also shows the highest performance metrics for classification accuracy (0.919), F1-score (0.919) and MCC values (0.838) indicating that it performs well on the human GENCODE dataset in comparison to other tools (Table 1A).

**Table 1A:** Tools performance on GENCODE human datasets. Bold-underlined values correspond the highest. CPAT_train and PLEK_train correspond to program versions trained with the HL dataset. Programs are sorted by MCC values.

| HUMAN dataset | Program | Sensitivity | Specificity | Precision | Accuracy | F1-score | MCC |
| --- | --- | --- | --- | --- | --- | --- | --- |
|  | FEELnc | **<u>0.923</u>** | 0.915 | 0.916 | **<u>0.919</u>** | **<u>0.919</u>** | **<u>0.838</u>** |
|  | CPAT | 0.899 | 0.924 | 0.922 | 0.912 | 0.91 | 0.823 |
|  | CPAT_train | 0.92 | 0.901 | 0.903 | 0.91 | 0.911 | 0.821 |
|  | CNCI | 0.829 | 0.979 | 0.975 | 0.904 | 0.896 | 0.817 |
|  | PLEK | 0.732 | **<u>0.985</u>** | **<u>0.981</u>** | 0.858 | 0.838 | 0.741 |
|  | PhyloCSF | 0.906 | 0.802 | 0.82 | 0.854 | 0.861 | 0.712 |
|  | PLEK_train | 0.582 | 0.96 | 0.936 | 0.77 | 0.718 | 0.584 |
|  | CPC | 0.699 | 0.739 | 0.728 | 0.719 | 0.713 | 0.438 |

We further investigated performance on the mouse datasets composed of 2,000 lncRNAs and 5,000 mRNAs in order to replicate the analysis in another organism and using an unbalanced dataset. For this benchmark, we included all the same programs except PhyloCSF due to the labour-intensive task to extract input cross-species multiple alignments. Again, FEELnc displays the highest classification accuracy, F-score and MCC (Table 1B) while CPC shows the best specificity (0.992) and precision (0.996) despite a weak sensitivity (0.744). As for human dataset, the CPAT program performs well even if we consider both the re-trained and prebuilt models. Interestingly, CPAT with the trained mouse model has only slightly better performance than using the human prebuilt model indicating that cross-species training achieves relatively few performance gains.

**Table 1B:** Program performances on GENCODE mouse datasets. Bold-underlined values correspond the highest. Programs are sorted by MCC values.

| MOUSE dataset | Program | Sensitivity | Specificity | Precision | Accuracy | F1-score | MCC |
| --- | --- | --- | --- | --- | --- | --- | --- |
|  | FEELnc | 0.938 | 0.941 | 0.976 | **<u>0.939</u>** | **<u>0.956</u>** | **<u>0.856</u>** |
|  | CPAT_train | **<u>0.95</u>** | 0.88 | 0.952 | 0.93 | 0.951 | 0.828 |
|  | CPAT | 0.892 | 0.96 | 0.982 | 0.911 | 0.935 | 0.806 |
|  | CNCI | 0.857 | 0.972 | 0.987 | 0.89 | 0.918 | 0.772 |
|  | CPC | 0.744 | **<u>0.992</u>** | **<u>0.996</u>** | 0.815 | 0.852 | 0.667 |
|  | PLEK | 0.71 | 0.913 | 0.954 | 0.768 | 0.814 | 0.564 |
|  | PLEK_train | 0.63 | 0.891 | 0.936 | 0.704 | 0.753 | 0.47 |

Finally, we assessed the computational time of each program including the time required for computing the model for trained-based tools. It results that FEELnc took ~46 min to classify the 10,000 human lncRNAs and mRNAs while PLEK was the fastest (6 min compared to ~10h when we train the model i.e. PLEK_train_) and CPC the longest (~2 days; with default parameters).

### 3. Annotating lncRNAs without a species-specific training set of lncRNAs

In the absence of species-specific lncRNAs set, machine-learning strategies require to simulate non-coding RNA sequences to train the model. As DNA composition varies between species, a first naive approach called *“intergenic”* consists in extracting random sequences from the genome of interest to model species-specific noncoding sequences. More precisely, using the human genome as example, the *intergenic* approach consists in randomly extracting *n* human sequences (with *n* corresponding to the number of input mRNAs) that do not overlap any mRNAs from the reference annotation. Another more sophisticated approach involves that lncRNAs derived from “debris” of protein-coding genes as exemplified by the *Xist* lncRNA that emerged from the disruption of the mRNA gene *Lnx3* (46). For this strategy that we called *“shuffle”,* we took the 5,000 HL mRNAs and shuffled the sequences while preserving *k*-mer frequencies using Ushuffle (39). We tested different size of *k* and showed that preserving 7-mer frequencies gave the best MCC values on the HT set while sustaining a high number of permuted sequences (Supplementary Fig.6). In order to evaluate the *intergenic* and *shuffle* strategies on the training sets, we computed their predictor scores in comparison with the true set of 5,000 HL lncRNAs. We first observed that the cumulative distribution of predictor values for the *shuffle* strategy tend to be closer to the one observed in the true set of human lncRNAs compared to the *intergenic* approach (Supplementary Fig.7). We then confirmed on the HT dataset that the *shuffle* method outperformed the *intergenic* approach (MCCs = 0.768 versus 0.646) as compared to true set of lncRNAs (MCC = 0.846) (Supplementary Fig.7).

As described above, FEELnc can be used in a stringent mode in order to distinguish and annotate high-confidence set of lncRNAs and mRNAs. To this end, the 2-threshold module was applied for both strategies with increasing specificity thresholds (0.93, 0.96, 0.99) as compared to the automatic optimal CPS cut-off defined previously. With these cut-offs, we observed higher performances for the *shuffle* approach (MCCs = 0.823, 0.881 and 0.943) versus *intergenic* (MCCs = 0.646, 0.654 and 0.785) (Supplementary Fig.7). Moreover, the greater interest of the *shuffle* approach could be appreciated by the lower variability between sensitivity and specificity metrics as defined within the FEELnc methodology compared to the *intergenic* approach (Supplementary Fig.7).

In order to directly evaluate FEELnc performance for “non-model” organisms, we used the FEELnc shuffle strategy where the coding parameters were learned on species-specific mRNAs and the noncoding ones on species-specific mRNAs shuffled by Ushuffle with preserved *7*-mer frequencies. All tests were further assessed on the catalogues of lncRNAs annotated in NONCONDE database. We also compared performances with CNCI knowing that NONCODE uses both CNCI and matching protein-coding coordinates from RefSeq database to remove all ncRNAs annotated as protein coding. In Supplementary Table 3, we showed that FEELnc without annotated lncRNAs achieved good classification metrics in many species (MCC ranging from 0.70 for rat to 0.903 for cow) and even better than CNCI in five species (nematode, fly, gorilla, orangutan and rhesus macaque).

As a third approach to model noncoding sequences, we sought to use cross-species lncRNAs for learning FEELnc model parameters. To this end, NONCODE lncRNAs from 13 organisms were used to serve as proxy for human lncRNAs and performance were then evaluated on the HT dataset. As expected, FEELnc performance is negatively correlated with the time of speciation between human and NONCODE species (spearman rho = −0.85; p-val = 5.6e−05) with MCC values of 0.374 when using *Caenorhabditis elegans* lncRNAs to 0.823 with chimpanzee lncRNAs (Supplementary Table 4). This probably reflects the variability in term of lncRNA sequence conservation (and thus in noncoding *k*-mer frequencies) between human and NONCODE species (35, 42). Interestingly, the *shuffle* strategy that has a MCC of 0.748 would correspond to performance obtained with lncRNAs that diverged about 100 Myr indicating that it constitutes an interesting approach when no lncRNAs from closely related species are available.

### 4. Application to identify an extended catalogue of canine lncRNAs

Within the framework of the LUPA consortium (29), we performed 20 whole transcriptome sequencing of 16 tissues (Supplementary Table 1). After QC, ~1,3 billions reads were mapped onto the canfam3 genome using the STAR mapper (47). Cufflinks (11) was further used to reconstruct the transcriptome models in each tissue separately guided by a consensus reference annotation given by the BROAD (48) and EnsEMBL v83 (22) annotations (called canFam3.1). Then, the cuffmerge tool merged the tissue samples files into a single GTF file containing 211,794 transcripts and 50,547 genes. In order to annotate new transcribed loci, we used the FEELnc_filter_ module to remove all transcripts which overlap exons (in sense) from the reference annotation, single exonic intergenic transcripts and transcripts with a size below 200 nt. A total of 5,523 candidate transcripts were retained and then given to FEELnc_codpot_ analysis. The two-thresholds option was run with a minimal specificity threshold fixed at 0.93 in both biotypes. This cut-off allows leveraging the number of ambiguous transcripts (TUCps) while optimizing classification specificity. This analysis identified 3,822 novel lncRNA transcripts, 477 new mRNA transcripts and 884 TUCps.

In a second analysis, we aimed at developing a comprehensive annotation of novel transcript isoforms based on RNA-seq assembled models overlapping (but not included in) canFam3.1 annotation (n = 69,602) (See Supplementary Methods for details). On this subset of assembled transcripts, we used FEELnc_codpot_ with the exact same parameters as previously. Because a novel transcript isoform could fuse two or more genes from the reference annotation, we defined rules to avoid the merging of transcripts having different biotypes by removing these incompatible transcripts (see Supplementary Methods for details). This resulted in the annotation of 67,312 transcripts with much more mRNA isoforms (58,163) compared to lncRNAs (6,552) and TUCps (2,597).

At the end, the new canine annotation that we called canFam3.2, includes a total of 36,237 loci with 3,145 new genes and 10,374 novel lncRNA, 58,640 new mRNA and 3,481 TUCp transcripts (Supplementary Table 1).

This study improves the dog genome annotation by several factors. First, we extended the number of mRNA transcripts by 50% and lncRNA transcripts by ~100%. Second, we found that novel lncRNA and mRNA transcripts are longer in term of number of exons, CDS/UTRs and RNA sizes compared to canFam3.1 transcripts suggesting a more complete reconstruction of their gene structures (Supplementary Table 6). Also, the number of isoforms per gene locus has been expanded to ~2.19 and ~7.2 for lncRNAs and mRNAs, respectively. Third, using STAR and RSEM (49) to quantify transcripts expression levels, we found that 86% of novel lncRNAs have at least a TPM (transcript per million) value >0.5 in at least one of the 20 samples highlighting a robust set of new lncRNAs. Finally, among the 2,658 canine novel lncRNA genes, 15% are also found in human GENCODE annotation by using the EnsEMBL compara EPO alignments (50). For instance, canFam3.2 now annotates three Cancer Susceptibility candidates lincRNAs (*CASC* lincRNAs) such as the *CASC9* lincRNA located on chr29:23,554,585-23,605,371 and involved in esophageal squamous cell carcinoma (51). Other examples include the well-described *MALAT1* cancer-associated lincRNA (52) which was considered as an unclassified non-coding transcript in canFam3.1 and the *IFNG-AS* antisense lncRNA involved in T-cell differentiation (53).

By employing the FEELnc_classifier_ on the new canFam3.2 transcripts, we identified 2,130 intergenic lncRNAs (lincRNAs) and 342 divergent lincRNAs closely located to their mRNA partner (distance < 5kb) highlighting potential bi-directional promoters. Conversely, FEELnc_classifier_ identified 276 antisense exonic lncRNAs and 308 sense exonic lncRNAs overlapping short ncRNAs. This canFam3.2 annotation constitutes a new resource that will help providing lncRNA as candidates for understanding and establishing genotype to phenotype relationships.

## DISCUSSION

In this study, we designed a new program to identify and annotate lncRNAs called FEELnc for FlExible Extraction of Long non-coding RNAs. Using the gold standard GENCODE annotation in human and mouse (27), we showed that FEELnc performs well to discriminate long non-coding versus protein-coding RNAs. FEELnc include predictors (multi *k*-mer frequencies and ORF coverage) that are general enough to capture all lncRNAs classes whereas alignment-based methods will be biased towards misclassifying species-specific transcripts or coding transcripts that are not referenced in peptide databases. The integration of a random forest-based model in FEELnc contributes to the good achievements of the tool because of the intrinsic properties of randomly sub-sampling features thereby encompassing diverse lncRNA characteristics. In recent years, major advances have been made in the machine-learning field (54) allowing to cope with the thousands of parameters simultaneously and it would thus be of great value to adapt deep learning approaches to coding and non-coding RNA annotations.

The contribution of FEELnc is not only limited to provide high classification performance metrics on models organisms since the tool is also accompanied with several modules and options that enable fine-tuning and precisely adjusting lncRNA annotations for any species of interest. To our knowledge, it is the first tool allowing users to annotate conservative sets of lncRNAs and mRNAs by fixing their own specificity thresholds(38). Second, FEELnc can be used for any given species even in the absence of lncRNA training set due to the possibility to model species-specific lncRNAs. Third, we expect the FEELnc classifier module to be of great interest to researchers in order to automatically annotate novel lncRNAs thus directly providing candidate pairs of lncRNAs and (m)RNA partners to be investigated for experimental validations.

For non-model organisms, we have shown that FEELnc performs similarly to CNCI although the benchmark was done on NONCODE lncRNAs that were *a priori* filtered by the CNCI tool. As for human and mouse where the manually curated GENCODE annotation is considered as a standard, this stressed the importance of also defining gold-standard sets of lncRNAs/mRNAs to correctly evaluate programs for “non-model” organisms annotation. This could be envisaged within the framework of collaborative projects such as the FAANG (30) for instance.

Finally, we illustrated the usefulness of FEELnc on the dog transcriptome for which 20 RNA-seq samples were sequenced in the frame of the LUPA consortium. The biological relevance of this expanded canine resource is supported by the annotation of novel dog lncRNAs orthologous to known lncRNAs in human and the increased transcripts and CDS sizes as well as exon numbers suggesting more complete RNA structures. Although this improved annotation will facilitate the identification of genotype to phenotype associations, it is still incomplete given the plethora of RNA classes beside lncRNA and mRNAs. For instance, one could consider annotating transcribed (processed or unprocessed) pseudogenes or enhancer RNAs (eRNAs) by using FEELnc as a multiclass classifier instead of a binary classifier (e.g coding versus noncoding). Some preliminary results on 1,123 human transcribed pseudogenes annotated in GENCODE showed that their FEELnc CPS distribution (mean CPS = 0.622) is clearly in-between lncRNAs and mRNAs CPS (mean CPS = 0.085 and 0.923, respectively) suggesting that it would be feasible to discriminate pseudogenes from lncRNA and mRNA. Similarly, eRNAs defined as non-coding transcripts derived from enhancer elements (55, 56) should harbor small ORFs and specific patterns of *k*-mers corresponding to the transcription factor binding sites which could be caught by FEELnc predictors.

All together, the FEELnc tool provides a standardized and exhaustive protocol to identify and annotate lncRNAs.

## ACKNOWLEDGEMENT

The authors are grateful to referring veterinarians, all dog owners and breeders for sample collection and also people from the European LUPA consortium (www.eurolupa.eu). We also thank FEELnc users and more particularly Kevin Muret, Sandrine Lagarrigue, Mark cock and Sarah Djebali. Finally, we are grateful to Jacques Nicolas for useful discussions and the GenOuest Bioinformatics core facility (www.genouest.org) for storing sequencing data, hosting the Cani-DNA website and for the use of the bioinformatic cluster to analyse the data.

## REFERENCES

1. Djebali, S., Davis, C.A., Dobin, A., Lassmann, T., Mortazavi, A., Tanzer, A., Lagarde, J., Lin, W., Schlesinger, F., Xue, C., et al. (2012) Landscape of transcription in human cells. Nature, 488, 101–108.

2. Pervouchine, D.D., Djebali, S., Breschi, A., Davis, C.A., Barja, P.P., Dobin, A., Tanzer, A., Lagarde J., Zaleski, C., See, L-H., et al. (2015) Enhanced transcriptome maps from multiple mouse tissues reveal evolutionary constraint in gene expression. Nat Commun, 6, 1–11.

3. Carninci, P., Kasukawa, T., Katayama, S., Gough, J., Frith, M.C., Maeda, N., Oyama, R., Ravasi, T., Lenhard, B., Wells, C., et al. (2005) The transcriptional landscape of the mammalian genome. Science, 309, 1559–1563.

4. Brown J.B., Boley, N., Eisman, R., May, G.E., Stoiber, M.H., Duff, M.O., Ben W Booth, Wen, J., Park, S., Suzuki, A.M., et al. (2014) Diversity and dynamics of the Drosophila transcriptome. Nature, 10.1038/nature12962.

5. Legeai, F. and Derrien, T. (2015) Science Direct Identification of long non-coding RNAs in insects genomes. Current Opinion in Insect Science, 7, 37–44.

6. Paytuví Gallart, A., Hermoso Pulido, A., Anzar Martínez de Lagrán, I., Sanseverino, W. and Aiese Cigliano, R. (2016) GREENC: a Wiki-based database of plant lncRNAs. Nucleic Acids Res, 44, D1161–6.

7. Li, L., Eichten, S.R., Shimizu, R., Petsch, K., Yeh, C.-T., Wu, W., Chettoor, A.M., Givan, S.A., Cole, R.A., Fowler, J.E., et al. (2014) Genome-wide discovery and characterization of maize long non-coding RNAs. Genome Biol, 15, R40–15.

8. Brockdorff, N., Ashworth, A., Kay, G.F., McCabe, V.M., Norris, D.P., Cooper, P.J., Swift, S. and Rastan, S. (1992) The product of the mouse Xist gene is a 15 kb inactive X-specific transcript containing no conserved ORF and located in the nucleus. Cell, 71, 515–526.

9. Prensner, J.R. and Chinnaiyan, A.M. (2011) The Emergence of lncRNAs in Cancer Biology. Cancer Discovery, 1, 391–407.

10. Leucci, E., Vendramin, R., Spinazzi, M., Laurette, P., Fiers, M., Wouters, J., Radaelli, E., Eyckerman, S., Leonelli, C., Vanderheyden, K., et al. (2016) Melanoma addiction to the long non-coding RNA SAMMSON. Nature, 531, 518–522.

11. Trapnell, C., Williams, B.A., Pertea, G., Mortazavi, A., Kwan, G., van Baren, M.J., Salzberg, S.L., Wold, B.J. and Pachter, L. (2010) Transcript assembly and quantification by RNA-Seq reveals unannotated transcripts and isoform switching during cell differentiation. Nature Biotechnology, 28, 511–515.

12. Pertea, M., Pertea, G.M., Antonescu, C.M., Chang, T.-C, Mendell, J.T. and Salzberg, S.L. (2015) StringTie enables improved reconstruction of a transcriptome from RNA-seq reads. Nat Biotechnol, 33, 290–295.

13. Haas, B.J., Papanicolaou, A., Yassour, M., Grabherr, M., Blood, P.D., Bowden, J., Couger, M.B., Eccles, D., Li, B., Lieber, M., et al. (2013) De novo transcript sequence reconstruction from RNA-seq using the Trinity platform for reference generation and analysis. Nat Protoc, 8, 1494–1512.

14. Lin, M.F., Jungreis, I. and Kellis, M. (2011) PhyloCSF: a comparative genomics method to distinguish protein coding and non-coding regions. Bioinformatics, 27, i275–i282.

15. Kong, L., Zhang, Y., Ye, Z.Q., Liu, X.Q., Zhao, S.Q., Wei, L. and Gao, G. (2007) CPC: assess the protein-coding potential of transcripts using sequence features and support vector machine. Nucleic Acids Res, 35, W345–W349.

16. Sun, L., Luo, H., Bu, D., Zhao, G., Yu, K., Zhang, C., Liu, Y., Chen, R. and Zhao, Y. (2013) Utilizing sequence intrinsic composition to classify protein-coding and long non-coding transcripts. Nucleic Acids Res, 41, e166.

17. Li, A., Zhang, J. and Zhou, Z. (2014) PLEK: a tool for predicting long non-coding RNAs and messenger RNAs based on an improved k-mer scheme. BMC Bioinformatics, 15, 1–10.

18. Wang, L., Park, H.J., Dasari, S., Wang, S., Kocher, J.-P. and Li, W. (2013) CPAT: Coding-Potential Assessment Tool using an alignment-free logistic regression model. Nucleic Acids Res, 41, e74–e74.

19. Dewey, C.N. and Pachter, L. (2006) Evolution at the nucleotide level: the problem of multiple whole-genome alignment. Human Molecular Genetics, 15 Spec No 1, R51–6.

20. Brent, M.R. (2007) How does eukaryotic gene prediction work? Nat Biotechnol, 25, 883–885.

21. Blanco, E., Parra, G. and Guigó, R. (2007) Using geneid to identify genes. Curr Protoc Bioinformatics, Chapter 4, Unit 4.3.

22. Yates, A., Akanni, W., Amode, M.R., Barrell, D., Billis, K., Carvalho-Silva, D., Cummins, C., Clapham, P., Fitzgerald, S., Gil, L., et al. (2016) Ensembl 2016. Nucleic Acids Res, 44, D710–6.

23. Johnson, R. and Guigo, R. (2014) The RIDL hypothesis: transposable elements as functional domains of long noncoding RNAs. RNA, 20, 959–976.

24. Zucchelli, S., Fasolo, F., Russo, R., Cimatti, L., Patrucco, L., Takahashi, H., Jones, M.H., Santoro, C., Sblattero, D., Cotella, D., et al. (2015) SINEUPs are modular antisense long non-coding RNAs that increase synthesis of target proteins in cells. Front. Cell. Neurosci., 9, 1720–1712.

25. Tagu, D., Colbourne, J.K. and gre, N.N. (2014) Genomic data integration for ecological and evolutionary traits in non-model organisms. 15, 1–16.

26. Breiman, L. (2001) Random Forests. Machine Learning, 45, 5–32.

27. Harrow, J., Frankish, A., Gonzalez, J.M., Tapanari, E., Diekhans, M., Kokocinski, F., Aken, B.L, Barrell, D., Zadissa, A., Searle, S., et al. (2012) GENCODE: The reference human genome annotation for The ENCODE Project. Genome Res, 22, 1760–1774.

28. Zhao, Y., Li, H., Fang, S., Kang, Y., Wu, W., Hao, Y., Li, Z., Bu, D., Sun, N., Zhang, M.Q., et al. (2016) NONCODE 2016: an informative and valuable data source of long non-coding RNAs. Nucleic Acids Res, 44, D203–8.

29. Lequarré, A.-S., Andersson, L., André, C., Fredholm, M., Hitte, C., Leeb, T., Lohi, H., Lindblad-Toh, K. and Georges, M. (2011) LUPA: A European initiative taking advantage of the canine genome architecture for unravelling complex disorders in both human and dogs. The Veterinary Journal, 189, 155–159.

30. Andersson, L., Archibald, A.L, Bottema, C.D., Brauning, R., Burgess, S.C, Burt, D.W., Casas, E., Cheng, H.H., Clarke, L., Couldrey, C., et al. (2015) Coordinated international action to accelerate genome-to-phenome with FAANG, the Functional Annotation of Animal Genomes project. Genome Biol, 16, 57.

31. Bussotti, G., Raineri, E., Erb, I., Zytnicki, M., Wilm, A., Beaudoing, E., Bucher, P. and Notredame, C. (2011) BlastR--fast and accurate database searches for non-coding RNAs. Nucleic Acids Res, 39, 6886–6895.

32. Drezen, E., Rizk, G., Chikhi, R., Deltel, C., Lemaitre, C., Peterlongo, P. and Lavenier, D. (2014) GATB: Genome Assembly & Analysis Tool Box. Bioinformatics, 30, 2959–2961.

33. Claverie J.M., Sauvaget, I. and Bougueleret, L. (1990) K-tuple frequency analysis: from intron/exon discrimination to T-cell epitope mapping. Meth. Enzymol., 183, 237–252.

34. Cabili, M.N., Trapnell, C., Goff, L., Koziol, M., Tazon-Vega, B., Regev, A. and Rinn J.L (2011) Integrative annotation of human large intergenic noncoding RNAs reveals global properties and specific subclasses. Genes & Development, 10.1101/gad.17446611.

35. Derrien, T., Johnson, R., Bussotti, G., Tanzer, A., Djebali, S., Tilgner, H., Guernec, G., Martin, D., Merkel, A., Knowles, D.G., et al. (2012) The GENCODE v7 catalog of human long noncoding RNAs: analysis of their gene structure, evolution, and expression. Genome Res, 22, 1775–1789.

36. Lertampaiporn, S., Thammarongtham, C., Nukoolkit, C., Kaewkamnerdpong, B. and Ruengjitchatchawalya, M. (2014) Identification of non-coding RNAs with a new composite feature in the Hybrid Random Forest Ensemble algorithm. Nucleic Acids Res, 10.1093/nar/gku325.

37. Sing, T., Sander, O., Beerenwinkel, N. and Lengauer, T. (2005) ROCR: visualizing classifier performance in R. Bioinformatics, 21, 3940–3941.

38. Mattick J.S. and Rinn, J.L. (2015) Discovery and annotation of long noncoding RNAs. Nature Structural & Molecular Biology, 22, 5–7.

39. Jiang, M., Anderson, J., Gillespie, J. and Mayne, M. (2008) uShuffle: A useful tool for shuffling biological sequences while preserving the k-let counts. BMC Bioinformatics, 9, 192–11.

40. Askarian-Amiri, M.E., Crawford, J., French, J.D., Smart, C.E., Smith, M.A., Clark, M.B., Ru, K., Mercer, T.R., Thompson, E.R., Lakhani, S.R., et al. (2011) SNORD-host RNA Zfas1 is a regulator of mammary development and a potential marker for breast cancer. RNA, 17, 878–891.

41. Lykke-Andersen, S., Chen, Y., Ardal, B.R., Lilje, B., Waage, J., Sandelin, A. and Jensen, T.H. (2014) Human nonsense-mediated RNA decay initiates widely by endonucleolysis and targets snoRNA host genes. Genes & Development, 28, 2498–2517.

42. Chen, J., Shishkin, A.A., Zhu, X., Kadri, S., Maza, I., Guttman, M., Hanna, J.H., Regev, A. and Garber, M. (2016) Evolutionary analysis across mammals reveals distinct classes of long non-coding RNAs. Genome Biol, 10.1186/s13059-016-0880-9.

43. Sigova, A.A., Mullen, A.C., Molinie, B., Gupta, S., Orlando, D.A., Guenther, M.G., Almada, A.E., Lin, C., Sharp, P.A., Giallourakis, C.C, et al. (2013) Divergent transcription of long noncoding RNA/mRNA gene pairs in embryonic stem cells. Proceedings of the National Academy of Sciences, 110, 2876–2881.

44. Carrieri, C., Cimatti, L., Biagioli, M., Beugnet, A., Zucchelli, S., Fedele, S., Pesce, E., Ferrer, I., Collavin, L., Santoro, C., et al. (2012) Long non-coding antisense RNA controls Uchl1 translation through an embedded SINEB2 repeat. Nature, 491, 454–457.

45. He, Y., Vogelstein, B., Velculescu, V.E., Papadopoulos, N. and Kinzler, K.W. (2008) The antisense transcriptomes of human cells. Science, 322, 1855–1857.

46. Duret, L., Chureau, C., Samain, S., Weissenbach, J. and Avner, P. (2006) The Xist RNA gene evolved in eutherians by pseudogenization of a protein-coding gene. Science, 312, 1653–1655.

47. Dobin, A., Davis, C.A., Schlesinger, F., Drenkow, J., Zaleski, C., Jha, S., Batut, P., Chaisson, M. and Gingeras, T.R. (2013) STAR: ultrafast universal RNA-seq aligner. Bioinformatics, 29, 15–21.

48. Hoeppner, M.P., Lundquist, A., Pirun, M., Meadows, J.R.S., Zamani, N., Johnson, J., Sundström, G., Cook, A., FitzGerald, M.G., Swofford, R., et al. (2014) An Improved Canine Genome and a Comprehensive Catalogue of Coding Genes and Non-Coding Transcripts. PLoS ONE, 9, e91172.

49. Li, B. and Dewey, C.N. (2011) RSEM: accurate transcript quantification from RNA-Seq data with or without a reference genome. BMC Bioinformatics, 12, 323.

50. Herrero, J., Muffato, M., Beal, K., Fitzgerald, S., Gordon, L., Pignatelli, M., Vilella, A.J., Searle, S.M.J., Amode, R., Brent, S., et al. (2016) Ensembl comparative genomics resources. Database (Oxford), 2016, bav096.

51. Hao, Y., Wu, W., Shi, F., Dalmolin, R.J., Yan, M., Tian, F., Chen, X., Chen, G. and Cao, W. (2015) Prediction of long noncoding RNA functions with co-expression network in esophageal squamous cell carcinoma. BMC Cancer, 15, 1.

52. Gutschner, T., Hämmerle, M., Eissmann, M., Hsu, J., Kim, Y., Hung, G., Revenko, A., Arun, G., Stentrup, M., Gross, M., et al. (2013) The noncoding RNA MALAT1 is a critical regulator of the metastasis phenotype of lung cancer cells. Cancer Research, 73, 1180–1189.

53. Hu, G., Tang, Q., Sharma, S., Yu, F., Escobar, T.M., Muljo, S.A., Zhu, J. and Zhao, K. (2013) Expression and regulation of intergenic long noncoding RNAs during T cell development and differentiation. Nature Immunology, 14, 1190–1198.

54. LeCun, Y., Bengio, Y. and Hinton, G. (2015) Deep learning. Nature, 521, 436–444.

55. ørom, U.A. and Shiekhattar, R. (2013) Long noncoding RNAs usher in a new era in the biology of enhancers. Cell, 154, 1190–1193.

56. Ren, B. (2010) Transcription: Enhancers make non-coding RNA. Nature, 465, 173–174.

57. Hedges, S.B., Marin, J., Suleski, M., Paymer, M. and Kumar, S. (2015) Tree of life reveals clock-like speciation and diversification. Mol Biol Evol, 32, 835–845.

